# Self-organized patterning of peristaltic waves by suppressive actions in a developing gut

**DOI:** 10.1101/2024.11.20.624500

**Authors:** Koji Kawamura, Soichiro Kato, Shota Utsunomiya, Yoshiko Takahashi, Masafumi Inaba

## Abstract

Gut peristalsis is a wave-like movement of a local contraction along the gut, and plays important roles in nutrient digestion and absorption. When peristaltic waves emerge in embryonic guts, randomly distributed origins of peristaltic waves (OPWs) become progressively confined to specific sites. We have investigated how this random-to-organized positioning is achieved using the caecum as a model in chicken embryos. While prominent OPWs, recognized as active (spontaneous) contractions, are located at endpoints of the intact caecum, other regions are also found to possess latent rhythm unveiled by fragmentation of a caecum into pieces, showing that the latent rhythm is normally suppressed in the intact gut. Analyses with caecum fragments demonstrate that the latent rhythm is spatially patterned in an early gut, to which negative impact by primitive passing waves contributes; the more passing waves a region experiences to undergo forced/passive contractions, the slower latent rhythm this region acquires. This patterned latent rhythm underlies the final positioning of OPWs at later stages, where a site with faster latent rhythm dominates neighboring slower rhythm, surviving as a “winner” by macroscopic lateral inhibition. Thus, the random-to-organized patterning of OPWs proceeds by self-organization within the caecum, in which two distinct mechanisms, at least, are employed; suppressive actions by primitive waves followed by macroscopic lateral inhibition.

## Introduction

Gut peristalsis is a series of wave-like movements of a local contraction along the gut axis. Peristaltic movements propel internal contents, and are essential for nutrient digestion and absorption. Dysregulation of peristalsis is associated with many intestinal disorders underscoring the importance of understanding the mechanisms underlying the peristaltic movements. Peristaltic movements have extensively been studied for adult guts, in which distinguishment between extrinsic effects (e.g., mechanical stimulation by ingested material) and intrinsic mechanisms has been a challenge. To circumvent these, the embryonic gut has emerged to serve as a powerful model since it undergoes peristaltic movements without ingested contents (Chevalier 2017, Shikaya 2022).

The gut of chicken embryos is particularly useful since the spatial information along the entire gut can be obtained due to spatial landmarks along the gut axis. We previously reported a distribution map of <u>o</u>rigins of <u>p</u>eristaltic <u>w</u>aves (OPWs), which are sites undergoing spontaneous contraction followed by wave propagation, in the entire gut posterior to the duodenum. The map demonstrated that OPWs are progressively confined to specific sites along the gut axis as development proceeds (Shikaya et al 2022).

In the present study, we asked how OPWs are confined to specific sites using an embryonic caecum as a model. Unlike mammals, most avians possess a pair of caeca that protrude in a lateral-to-anterior direction from the main tract at the border between midgut (ileum) and hindgut (Fig. 1A). In chickens, left and right caeca elongate in parallel as straight tubular structures. These characteristics offer advantages; one part of caeca is used for experimental manipulation with the other being for control, minimizing complexities of variations between embryos. In addition, a short and straight shape facilitates quantitative assessment of OPW locations along the tubular structure, which is a challenge with a long and convoluted midgut. We demonstrate that while prominent OPWs are seen in the terminal endpoints of the caecum, whereas the middle region of the caecum displays less. However, when the caecum is fragmented into several small pieces along the gut axis, every piece commences to exhibit an OPW activity (spontaneous contraction) with their own contraction rhythm, unveiling latent rhythm possessed by every region along the caecum. Short-term (4 hours) monitoring shows that among subtle differences in latent rhythm along the caecum axis, a region with faster clock survives as a “winner” (OPW) by suppressing its neighbors’ rhythm of slower clock. In addition, a long-term study (3-day culture) reveals that the more frequently a given region experiences passing waves coming from OPWs, the slower rhythm this region acquires. Our findings provide evidence that patterning/positioning of OPWs along the gut is specified, at least in part, by two distinct types of self-organization mechanisms within the gut, long-term suppressive actions by passing waves followed by short-term macroscopic lateral inhibition.

**Figure 1.**
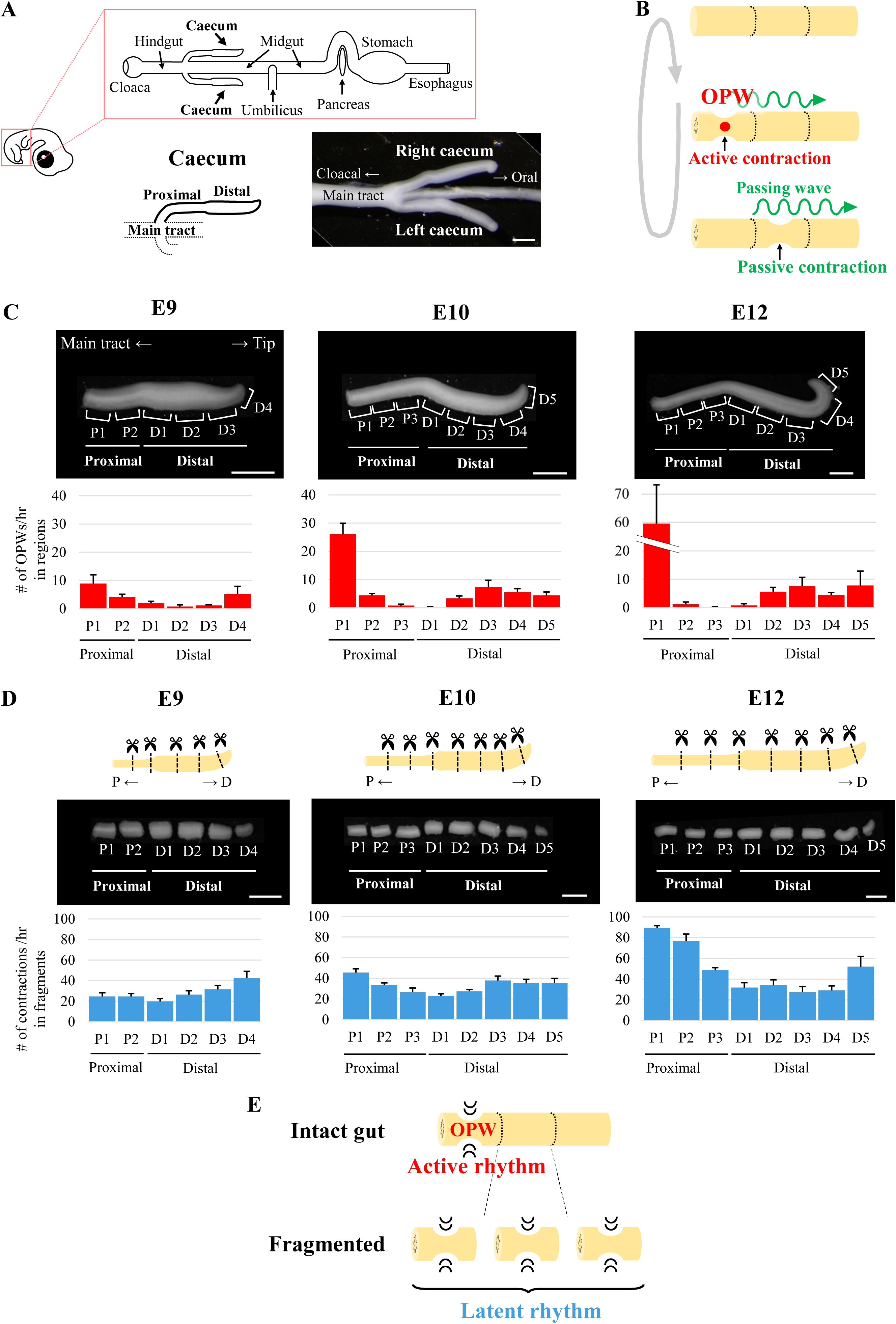
Spatial patterning of OPWs and latent rhythms during chicken caecum development. (**A**) Schematic representation of the chicken embryonic digestive tract (upper panel). Magnified view of the E10 caeca, shown as symmetrical protrusions from the main tract (lower panels). Scale bar, 1 mm. (**B**) Two types of gut contractions: 1) Active contraction: a spontaneous contraction at the site of OPW generation (red circle); 2) Passive contraction: a contraction induced by a passing wave (green wavy lines). (**C**, **D**) Developmental changes in the number of OPW generations per hour in each region (**C**) and the number of contractions per hour in each fragment (**D**) of the caecum. P, proximal; D, distal. Bar graphs and error bars represent average and standard error, respectively (n = 5 for each embryonic stage). Scale bars, 1 mm. (**E**) Summary diagram of Figure 1. In an intact gut, a specific region exhibits an active rhythm characterized by OPW generation. Upon separation, each fragment initiates its own distinct latent rhythm.

## Materials and Methods

### Chicken Embryos

Fertilized chicken eggs were obtained from the Yamagishi poultry farms (Wakayama, Japan), and embryos were staged according to Hamburger and Hamilton (HH) series (Hamburger and Hamilton, 1951) or embryonic day (E). All animal experiments were conducted with the ethical approval of Kyoto University (#202408).

### Short-term ex-vivo culture

Most experiments for short-term observation of gut motility (approximately less than 5 hours) were performed using the following culture conditions: Caeca were dissected from E9, E10, and E12 embryos and placed in a silicone-coated Petri dish (6 cm diameter) containing 13-15 ml of high-glucose DMEM (Wako, 048-29785) supplemented with 1× Penicillin-Streptomycin-Amphotericin B Suspension (PSA) (FUJIFILM, 161-23181). The Petri dish was maintained in a heating chamber (BLAST, C-140T and BE-051A) at 36.0°C under an atmosphere of 5% CO2 and 95% air. Six (proximal-1 to 2 and distal-1 to 4) and eight (proximal-1 to 3 and distal-1 to 5) equal-sized regions were defined in E9 caecum and E10, 12 caeca, respectively (Fig. 1C). Intact caeca (separated from the main tract) were imaged under free-moving conditions to minimize mechanical stress (Fig. 1C and Fig. 3A). In contrast, fragmented caeca based on each region (Fig. 1D, Fig. 3B lower, Fig. 4D right, and Fig. 4E right) were imaged within PDMS canals (internal dimensions: 0.8 mm x 1 mm) to prevent drift. Caeca were cultured for 2-4 hours until irregular gut movements subsided. Time-lapse images were acquired every 1-second for 1 hour using a Leica MZ10 F microscope equipped with a DS-Ri1 camera (Nikon). Image analysis of each region/fragment was performed using NIS Elements ER (Nikon, version 5.21.00). Images were processed and converted to intensity values, and the number of contractions was manually counted with the aid of image analysis.

### Provocation of the gut fragment contraction

The proximal 2 regions (P2) of E12 caeca were prepared as described above (Short-term ex-vivo culture). Caecal fragment was placed in a 3D-printed holder (made by Prusa i3 MK3S+) and fixed in a culture dish. To induce artificial gut contractions, a homemade mechanical stimulator was employed. A tungsten wire (Nilaco, 461167, 0.10 mm diameter) attached to a glass capillary was connected to a piezo bending actuator (THORLABS, PB4NB2W) via 3D-printed adaptors. The piezo actuator was controlled by a function waveform generator (Siglent, SDG1032X) via piezo driver (MESS-TEK, M-2691) with the following settings: Period, 1000 s; Amplitude, 3.000 Vpp; Offset, 0.000 Vdc; Pulse Width, 200.000 ms; Rise Edge, 16.8 ns; and Delay, 10 ms. This configuration ensured that the tungsten wire tapped the gut fragment immediately upon the movement of the actuator. The position of piezo bender was carefully adjusted with micromanipulator (Narishige, M-152) to tap appropriate region and avoid damaging the caecum fragment.

### Long-term ex-vivo culture

For 3-day incubation experiments (Fig. 3B upper panel, Fig. 4C, Fig. 4D left panel, and Fig. 4E left panel), caeca were dissected from E10 chicken embryos and placed in PDMS canals (internal dimensions: 0.7 mm x 1 mm) attached to a 6 cm diameter Petri dish containing 15 ml of high-glucose DMEM (Wako, 048-29785). The Petri dish was maintained in a heating chamber (BLAST, C-140T and BE-051A) at 38.5°C under 95% O2 and 5% CO2. Oxygen was supplied by an oxygen generator (KMC, M1O2S10L). For experiments shown in Fig. 3B upper panels, one caecum was randomly selected, and its proximal-1 (P1) and -2 (P2) regions were removed. The other caecum was left intact. For experiments shown in Fig. 4D left and 4E left panels, the P1 and P2 regions were removed from both caeca. Data from paired caeca were analyzed as control and intervention conditions. Time-lapse images were acquired every 1-second for 3 days, and contractions of distal-2 (D2) region were converted to intensity values using NIS Elements ER. A custom Python script (see Data analysis) was used to automatically detect passing waves as intensity peaks. In cases where automatic detection was unreliable, the number of contractions was manually counted.

### *In ovo* electroporation

*In ovo* electroporation was performed as previously described (Momose et al., 1999; Atsuta et al., 2013) with slight modifications. Briefly, a DNA solution was prepared at a concentration of 10 μg/μL, containing pT2A-CAGGS-ChR2(D156C)-mCherry-IRES-Neor: pCAGGS-T2TP: 4% fast green FCF (Wako, CI 42053) at a ratio of 4:1:0.5. This solution was injected into the coelomic cavity of the lateral plate mesoderm on the right side of Hensen’s node (presumptive right caecum) of E2.5 (HH13-15) stage embryos (Shikaya et al., 2023). An electrical pulse of 50 V, 0.05 ms duration was applied, followed by five pulses of 7 V, 25 ms duration, at 250 ms intervals (BEX, Pulse generator CUY21EDITⅡ). Electroporated embryos were incubated *in ovo* at 38.5°C with high humidity until E10 stage. Fluorescent images were acquired using a Leica MZ10 F microscope equipped with a DS-Ri1 camera (Nikon)

### Optogenetics

A pair of caeca, the right one expressing CRh2(D156C)-mCherry, was obtained from the E10 electroporated embryo described above. Right caecum expressing ChR2(D156C) served as the experimental group, while left caecum served as an internal control. Both caeca were dissected from the main tract, and the proximal-1 (P1) and -2 (P2) regions were removed. The caeca were then incubated using the previously described "Long-term ex-vivo culture" method. Blue light (λ = 470 nm) was generated by an LED (OSB56L5111P) and delivered focally to the distal-3 (D3) and -4 (D4) regions of the right (electroporated) caecum via a 500 μm diameter optical fiber. The fiber was inserted through a small hole in the lid of the Petri dish and positioned close to, but not in contact with, the specimen. Light pulse irradiation (40 ms pulse width, every 30 seconds) was controlled by a microcomputer Arduino Uno (arduino.cc). Irradiation typically induced the generation of OPWs at the irradiated region. These OPWs then propagated to the distal-2 (D2) region, observed as passing waves. Time-lapse images were acquired under red light (λ = 590 nm, Opto Code, EX-590 and LED-EXTA) for 3 days to minimize unintended activation of contractions. The number of passing waves in the D2 region was visually counted with the aid of NIS Elements ER analysis software. The number of contractions (times/hour) was calculated for the passing waves observed during organ culture. After 3 days of incubation, the D2 region was isolated from the specimens under red-light conditions. Time-lapse recordings were then performed to assess the number of contractions (times/hour) in the isolated region under "Short-term ex vivo Culture" conditions.

### Data analysis

Statistical analysis and graph construction were performed using Microsoft Excel. Paired t-tests were used to statistically compare the data presented in Fig. 3 and 4. Statistical significance was defined as *P < 0.05 and NS indicating not significant. Bar graphs and error bars represent the means and standard errors, respectively. Peaks associated with gut contractions were automatically detected using a SciPy library (scipy.signal.detrend, standardize, and find peaks) in Python. In cases where the automated detection was insufficient, peaks were manually counted.

## Results

### Spatial patterning of OPWs during caecum development

In this study, we define terms as follows: a spontaneous contraction at the site of OPW is called active contraction, and a contraction on the way of passing (propagating) wave is a passive contraction (Fig. 1B). Elongating caeca at E9 to E12 display a landmark at the middle along the proximal-to-distal (PD) axis; the proximal and distal portions are thinner (small in diameter) and thicker (large in diameter), respectively (Fig. 1A).

Caeca from E9, E10 and E12 embryos were excised from the main tract, and subjected to video monitoring for OPWs using an *ex vivo* culture method (see Materials and Methods). To precisely locate OPWs along the caecal axis, we divided the proximal and distal portions further into several regions (Fig. 1C, Movie S1, S2). The number of contractions per hour (frequency) in each region is shown in Fig. 1C. From E9 to E12 examined, originally indifferent frequency became differentiated between regions. For example, the proximal 1 region (P1) exhibited an increase in frequency during the time of development; 8.9 (E9), 26 (E10), and 59.6 (E12), whereas the frequency decreased in region P2; 4.1 (E9), 4.4 (E10), and 1.2 (E12). Of note, P2, P3 and D1 displayed no or few OPWs, if any, at E12 (Fig. 1C). Thus, the difference in OPW frequency among regions became enhanced indicating that OPWs undergo spatial patterning during caecal development.

### Latent rhythm was unveiled in non-OPW regions when isolated from neighbors

How are OPWs confined to specific regions in the caecum? Two possibilities are raised; one is that non-OPW regions (P2, P3, D1) have lost their contraction ability. Alternatively, contraction abilities are retained in these regions but somehow suppressed. To distinguish between these two, the caecum was cut into pieces to obtain fragments of P1 to D5, and assessed for contractions in each piece. Markedly, non-OPW fragments (P2, P3, D1) exhibited active contractions (Movie S3, Fig. 1D). The frequency in fragment P1, which already showed the highest in unfragmented (intact) caecum (Fig. 1C), was higher than fragment P2, and the same holds true for D5 (higher) and D4 (slower). Such differences between juxtaposed fragments (e.g., P1/Ps, D4/D5) were more pronounced at E12. In addition, when P1 was removed from an intact caecum at E12, P2 exhibited a markedly enhanced rhythm (1.8 to 35.6) (Fig. S1).

These findings suggest that: 1) every region in the cecum possesses the ability to generate active (spontaneous) contractions, hereafter called “latent rhythm”, and 2) the latent rhythm in non-OPW regions (P2, P3, D1) is suppressed by neighboring high-OPW regions.

### The latent rhythm is reset by interrupted contraction

Given that fragments P2-D1, which are normally non-OPW-generating regions in the intact caecum, exhibit latent rhythm upon isolation, we hypothesized that passing waves coming from a neighboring high rhythm OPW (e.g., P1) might mask the latent rhythm in intact caecum. To understand how this masking is regulated, we examined a reaction of a non-OPW-generating fragment (for example P2) to an ectopically induced contraction (interruption) that mimics a passive contraction forced to occur by a passing wave. Since a normal interval between spontaneous contractions (latent rhythm) of P2 at E12 was around 50 sec, we gave a mechano-stimulation by tapping at 40 sec after a first spontaneous contraction. A tapping with a tungsten wire controlled by a piezo actuator evoked contractions in fragment P2 in a consistent manner (Fig. 2A, Movie S4).

**Figure 2.**
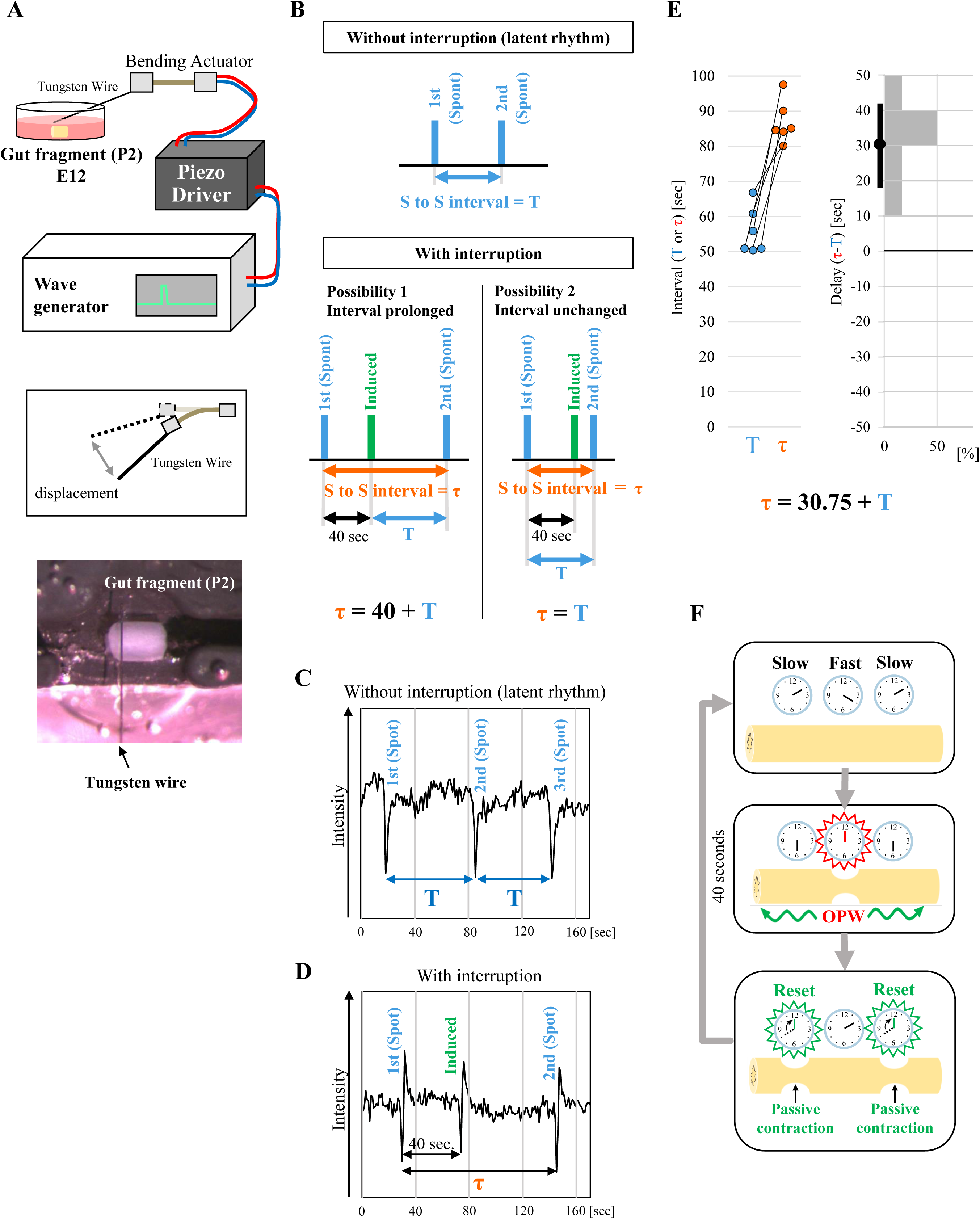
Resetting of the latent rhythm by an ectopically induced contraction. (**A**) Schematic of the mechanical stimulator setup. An E12 gut fragment (P2) in ex vivo culture was tapped by a thin tungsten wire attached to piezo bending actuator. The actuator was controlled by a wave generator via a driver (see Materials and Methods). The bottom panel shows an actual image of the tapping setup. (**B**) Expected results of rhythm interruption. Upper panel indicates the latent rhythm of P2 fragment without interruption. The interval between first and second spontaneous contractions (blue bars) is defined T. Lower panels indicate two possible outcomes with interruption. Upon an induced contraction by tapping (green bars), the interval between the first and second spontaneous contractions after the interruption (τ) can either be prolonged (τ = 40 seconds + T) or remain unchanged (τ = T). Representative data of temporal changes in P2 contractions without (**C**) and with interruption (**D**). Rapid changes in the traces represent spontaneous or induced contractions. (**E**) Quantification of the effects of interruption. The left graph shows the spontaneous interval without interruption (T) and with interruption (prolonged interval (τ)). Pairs of dots connected with a line represent the same specimen (n = 6). The right histogram shows the delay (difference between τ and T) in the latent rhythm caused by the interruption. The average of the delay and 95% confidence interval are shown as black dot and bar, respectively. (**F**) A model for rhythm resetting. Clocks represent the frequencies of latent rhythms. A faster clock in the middle generates OPWs that propagate to neighboring slower clocks and reset their rhythms. This mechanism allows the faster clock (higher latent rhythm) to maintain its rhythm and suppress the emergence of OPWs in slower clocks (slower latent rhythm).

Two possibilities for the P2 reaction were conceived. 1) The interval of latent rhythm would be delayed by the interruption; τ = 40 + T (T = normal interval of latent rhythm in blue, Fig. 2B, τ = total time (orange) between the first and second spontaneous contractions (blue)). 2) The latent rhythm would not be affected by the interruption; τ = T (Fig. 2B). Monitoring analyses showed that with a tapping interruption, the second spontaneous contraction took place at 86.83 sec after the first one (Fig. 2C, D, Movie S4 and Fig. 2E for paired comparison). Thus, τ = T + 30.75 ± 4.40 (average ± standard error), a result that is closer to the first possibility, that the interval of latent activity is delayed by the interruption.

With these findings, we propose a model as shown in Fig. 2F. Originally, every region in the embryonic caecum possesses its own “clock” of periodic contractions, which is slightly diverse between each other along the caecum axis. Among them, a region with faster clock emits propagating waves that force “passive” contractions in neighboring regions that have slower clock. Subsequently, the forced contracted regions “reset” their clock to count from zero. In this way, waves coming from a faster(est) clock defined as an OPW(s) continuously force passive contractions in neighboring slow regions, whose latent rhythm remains masked. We call these suppressive processes as “macroscopic lateral inhibition” in the gut, which is likely mediated by oscillatory entrainment known in physiology and neurobiology (Diamant and Rose, 1970, Van der Pol 1929, Perkel 1964).

### Latent rhythm was augmented with a reduced number of passing waves

Given that the spatial differences in latent rhythm (Fig. 2D) are the basis for self-organized sharp patterning (Fig. 2C) through macroscopic lateral inhibition (Fig. 2), we further explored how the differences in latent rhythm are generated at earlier stages of gut development. Because a non-OPW region is forced to contract (passive contraction) by an OPW-derived wave that passes over this region (passing waves), and also because such forced contractions reset the clock (Fig. 2), we reasoned that the differences in frequency of passing waves that are experienced by each region plays a role in shaping the differential patterns of latent rhythm along the caecum.

To address this, we examined a range of passing waves emitted from different sites of OPWs at E10. For example, as already shown in Fig. 1C, many waves were emitted from P1, and a substantial number of these waves vanished near D3 with a few propagating to the end of the caecum (Movie S1 and Fig. 3A; over 4.6 regions in average). Similarly, waves originated from other regions, although their frequencies were low, also ceased after propagating over 2 or 3 regions (Fig. 3A). Thus, the number of passing waves that each region experiences varied between regions; the middle regions experienced more waves, while the number of waves decreased toward both ends of the caecum. These differential patterns of passing wave experiences along the caecum are complementary with those shown for the latent rhythm (compare the graph in Fig. 3A with Fig. 1D).

**Figure 3.**
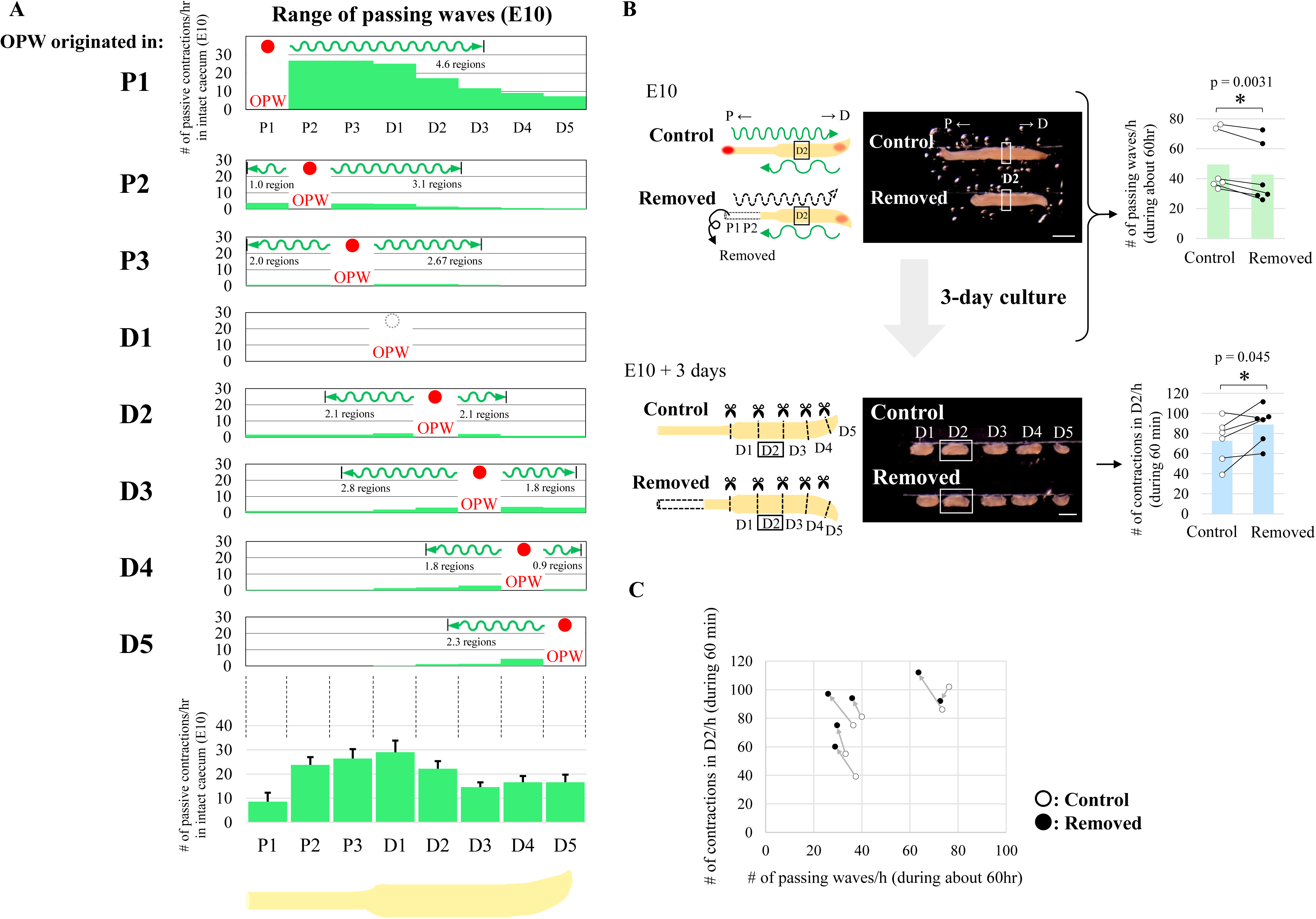
Reduction of passing waves accelerates latent rhythm. (**A**) Propagation range of passing waves in an intact E10 caecum. The upper panels show the number of passive contractions derived from passing waves of the OPW (red circles) in each region. The lower panel shows total number of passing waves for 1 hour observation in each region. (**B**) Long-term (3 days) ex vivo culture of E10 caeca with and without P1-P2 regions. Removal of the P1 and P2 regions reduced the number of passing waves per hour (upper panels). After 3 days of culture, the number of contractions in the D2 fragment increased compared to the control (lower panels). Bar graphs are represented as mean values. Paired data (same embryos) are connected by lines (n = 6 pairs). Scale bars, 1 mm. (**C**) Relationship between the number of passing waves and the acquired number of contractions in the D2 region of individual specimens.

To further comprehend how the passing wave experiences impact the production of latent rhythm, we attempted to reduce the frequency (number) of passing waves. When [P1-P2] was removed from a caecum, [P1-P2]-originated high frequent waves were extinguished leaving less frequent waves coming from distal tip [D4-5] unaffected (Fig. 3B, assessed for D2). The [P1-P2]-deprived specimens were *ex vivo* cultured for 3 days, and subjected to assessment of latent rhythm in fragments in a way similar to Fig. 1D. Contraction frequency of fragmented D2 was augmented compared with control (n=6, Fig. 3B in light blue), which was further corroborated by paired scatter plot shown in Fig. 3C. These findings imply that the passing waves negatively regulates the latent rhythm.

### Optogenetically augmented frequency of passing waves declined the latent rhythm

To more precisely scrutinize the role of passing waves in the differential distribution of latent rhythm, we experimentally increased the frequency of passing waves in the caecum using optogenetic methods that we recently developed for embryonic guts (Shikaya et al 2023). A modified type of channelrhodopsin 2, ChR2(D156C), efficiently responds to blue light irradiation to transmit depolarization signal intercellularly. Tol2-ChR2(D156C) tagged with mCherry was electroporated into the splanchnopleural mesoderm at the presumptive caecum region in the right half of embryo at E2 (Fig. 4A). ChR2(D156C)-expressing right- and untreated left caeca were dissected from embryos at E10 (Fig. 4B), followed by a removal of the proximal region [P1-P2] to obtain a less frequently waving caecum as shown above (Fig. 3). In these analyses, we irradiated blue light into D3/D4 at 30 sec intervals, which successfully evoked peristaltic waves passing at least over D2 with the rhythm obeying the irradiation cycle (Fig. 4C, Movie S5).

**Figure 4.**
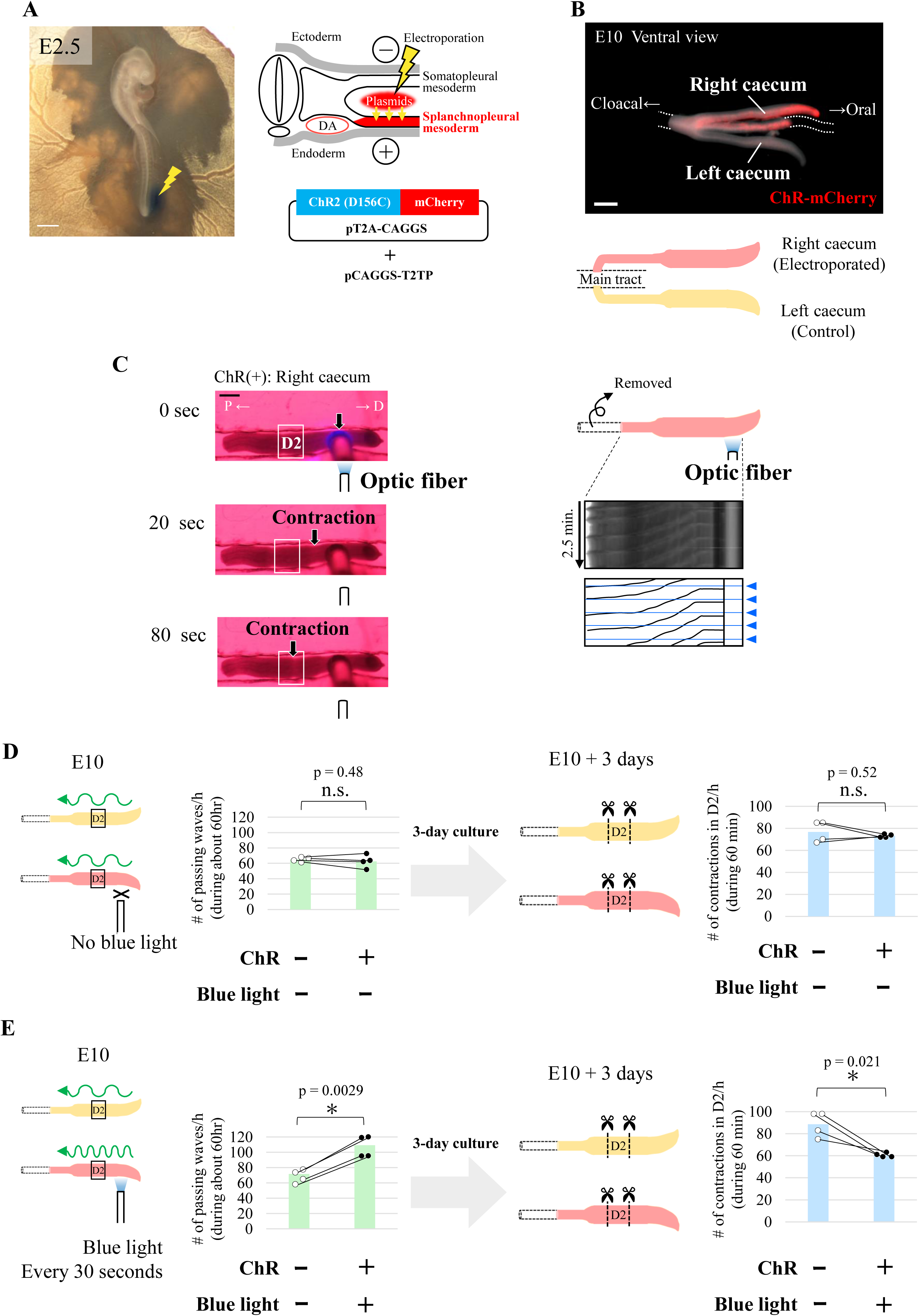
Optogenetically-increased passing waves decelerate latent rhythm. (**A**) *In ovo* electroporation of a Tol2 plasmid encoding channelrhodopsin-2 variant, ChR2(D156C), into the splanchnopleural mesoderm of the right side of the embryo at E2.5 (indicated by yellow zigzag lines). DA, dorsal aorta. Scale bar, 1 mm. (**B**) E10 caeca expressing ChR2(D156C) in the right caecum and a part of the main tract (dotted lines). The left caecum served as an internal control. Scale bar, 1 mm. (**C**) Left panels: Time-series images showing artificially induced peristaltic waves in an E10 caecum expressing ChR2(D156C). The P1 and P2 regions were removed. Blue light stimulation of the distal region triggered contractions that propagated to the D2 region (white rectangle). P, proximal; D, distal. Scale bar, 500 µm. Right panels: Kymographs showing serial peristaltic waves (black slanted lines) induced by 30-second intervals of blue light pulses (blue triangles). (**D, E**) Long-term ex-vivo culture of ChR2-D156C-expressing caeca without (**D**) and with (**E**) blue light stimulation. The P1 and P2 regions were removed. Green bars represent the frequency of passing waves through the D2 region during the 3-day culture. Blue bars represent the latent rhythm of the D2 fragment after 3 days of culture. Data are represented as mean values. Paired data (same embryos) are connected by lines (n = 4 pairs).

In ChR2(D156C)-expressing right caecum *without* blue light irradiation, contractions in D2 in both intact caecum and fragment were comparable to those in non-electroporated control (left caecum), showing no detectable effects by ChR2(D156C) overexpression (Fig. 4D). In contrast, when blue light was irradiated into D3/D4 every 30 seconds for 3 days, the number of passing waves were accordingly augmented. Importantly, the latent rhythm in fragment D2 prepared from the irradiated 3-day cultured caecum was significantly reduced (Fig. 4E). Thus, the more frequently a given region experiences passing waves, the slower rhythm it acquires (Fig. S2).

## Discussion

We have demonstrated that even non-OPW regions in the embryonic caecum retain an ability to implement active contraction unveiled by fragmentation of the gut. This ability is called latent rhythm in this study, which is normally suppressed in the intact gut. The latent rhythm becomes progressively patterned along the gut, and this patterning serves as the basis for the final confinement of OPWs at later stages. The patterning of OPWs proceeds by self-organization within the gut, where at least two distinct mechanisms are employed (Fig. 5). Step 1 is that initially less diverse latent rhythm among regions becomes more diverse/differentiated by the effects of randomly occurring passing waves; the more passing waves a given region experiences to undergo forced/passive contractions, the less frequent rhythm this region acquires. Step 2 is that a region with faster clock of latent rhythm that is generated by Step 1 dominates neighboring slow clock to survive as a winner. Step 2 is regarded as a process of macroscopic lateral inhibition mediated by entrainment. Both Step 1 and Step 2 employ suppressive actions of long-term (3 days) and short term (< one minute), respectively.

**Figure 5.**
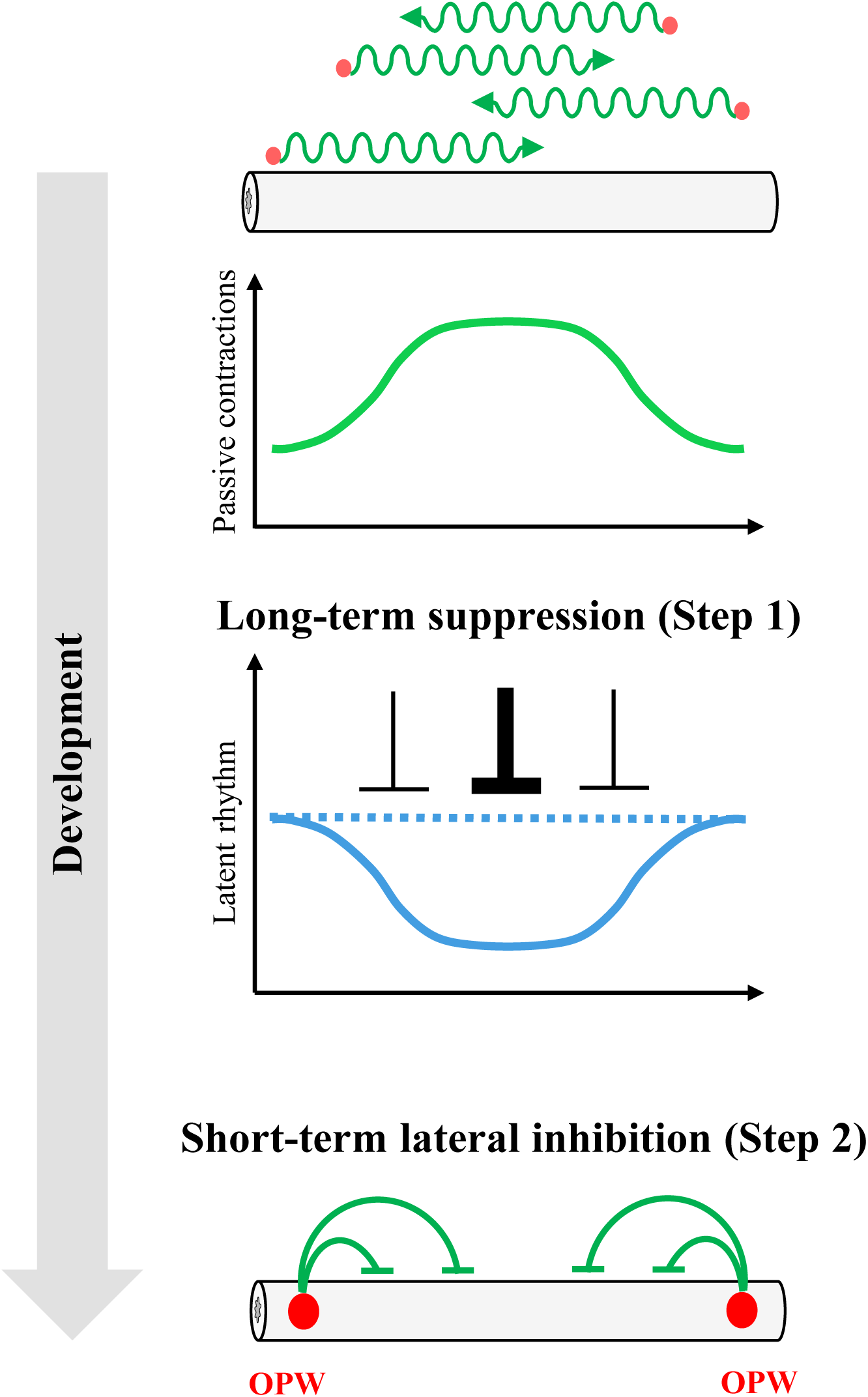
A model to explain progressive OPW patterning in developing gut. During gut development, passing waves (green wavy lines) emerge spontaneously at unspecified sites (red circles) along the gut axis. However, fluctuations of emergence of passing waves forms spatial gradient of passing wave frequency (bell-shaped green line) (Fig. 3A). This spatial gradient of passing wave frequency shapes a complementary spatial gradient of latent rhythms through long-term suppression (Fig. 3B, 3C and Fig. 4). The spatial gradient of latent rhythms ultimately specifies the location of the OPW region through a macroscopic lateral inhibition mechanism (green lines) (Fig. 2).

### Roles of precocious waves in the patterning of latent rhythm

While it is well known that peristaltic movements are important for the transportation of internal contents in adults, other roles of these movements remained poorly explored. In the current study using embryonic gut, we demonstrate that frequencies of peristaltic passing waves regulate negatively the latent rhythm. In [P1-P2]-lacking caecum showing a reduced number of passing waves, the latent rhythm was augmented after 3 days (Fig. 3). Conversely, when more passing waves were experimentally provided, the latent rhythm was declined. In early embryonic guts including caecum, primitive peristalsis-like movements occur randomly, where many waves vanish before reaching to the end (Chevalier 2017, Shikaya 2022 and this study), but the roles of these primitive/precocious waves have not been well understood. Our findings suggest that these waves in early guts negatively regulate the latent rhythm; the more frequently a given region experiences passing waves to undergo forced contractions, the slower latent rhythm it acquires. These negative regulations generate differential distribution of latent rhythm along the gut axis, which later shapes spatial patterns of OPWs by macroscopic lateral inhibition (see also below). The evidence to explain the regulation of OPW positioning by precocious peristaltic waves is unprecedented. How passing waves negatively impact on latent rhythm has yet to be studied. It was previously reported that reiterative stimulations by stretching cardiomyocyte *in vitro* resulted in converting its rhythm (Nitsan 2016), although this case was appositive correlation contrasting with our findings with a gut specimen.

### Macroscopic lateral inhibition of latent rhythm shapes the OPW positioning in embryonic gut

We have also found that in developing guts the differentiated patterns of latent rhythm generated by precocious passing waves (Step 1) is further subjected to macroscopic lateral inhibition, in which a region with faster-counting rhythm dominates its neighbors’ slow-counting regions (Step 2). A region having slower latent rhythm needs to obey faster rhythm to undergo forced/passive contraction without an opportunity to exert its own latent rhythm of active contraction. In such actions, also known as entrainment, a slow-counting region resets its rhythm when encountered with a faster rhythm (Fig. 2). If freed from entrainment, the latent rhythm can quickly be retrieved (as seen for augmented contractions in P2 in P1-lacking caecum, Fig. S1). The macroscopic lateral inhibition described in the current study is conceptually reminiscent of widely known lateral inhibition acting at the microscopic level, where Notch/Delta signaling mediates. In both micro- and macroscopic levels, initially equal properties become differentiated by mutual suppressive actions. Considering organs that are very long such as gut, long-range signals must be at work to coordinate patterning and growth, and it is sensible that the macroscopic lateral inhibition demonstrated in this study is one of the mechanisms that explain such organ-level coordination.

Latent rhythm (or its equivalent phenomena) in the gut was previously reported using adult intestine in felines (Diamant & Bortoff, 1969), where they proposed that linearly graded latent rhythm yields distinct sharp boundaries between contraction and non-contraction areas. More recently, a study using felines showed that when a region with fast rhythm in the intestine was occluded, a neighboring region started to exert a contraction rhythm, although slower than the occluded area (Lammers, Wim J 2008). Thus, the macroscopic lateral inhibition that we described in the current study could be a general mechanism underlying the patterning of peristaltic movements in the gut. Unlike studies with adult guts, our study addressed a transition from initially random patterns of early peristaltic movements to confined positioning at later stages, where macroscopic lateral inhibition (Step 2) is based on precocious conditions of passing waves (Step 1).

Regarding the entrainment in biological rhythm, similar phenomena have been described in cardiac pacemaker physiology. When the primary pacemaker of sinoatrial (SA) node becomes defective, latent rhythm of atrioventricular (AV) node, the second pacemaker, is retrieved and compensates for the failure of SA, although its rhythm is slower than that of SA (Alfred Zahn 1913, Robert P. Grant 1956, A L Waldo 1984, Leo Schamroth 1988). These notions suggest a similarity between heart and gut, and might help unveiling the molecular and cellular mechanisms underlying OPW-positioning in the gut.

## Acknowledgments

We thank Prof. Scott Gilbert for careful reading of the manuscript and discussion. We also thank Prof. Shinya Kawaguchi and Dr. Takuma Inoshita for technical advice for the oxygen generator, and National BioResource Project (Chicken-Quail, Nagoya University) for their technical help.

## Ethics statement

The animal study was reviewed and approved by Ethics committee for Regulations on Animal Experimentation at Kyoto University.

## Author contributions

KK, YT and MI conceived the study and wrote the manuscript. KK and MI performed experimental research. SK developed the mechanical stimulator. SU analyzed the data. All authors contributed to the article and approved the submitted version.

## Funding

This work was supported by JSPS KAKENHI Grant Numbers: 21K06198 (MI), 23H04702 (MI), 23H04933 (YT) and 23K19369 (SK); and by FY 2022 Kusunoki 125 of Kyoto University 125th Anniversary Fund (MI).

## Supplementary Figures

**Figure S1.**
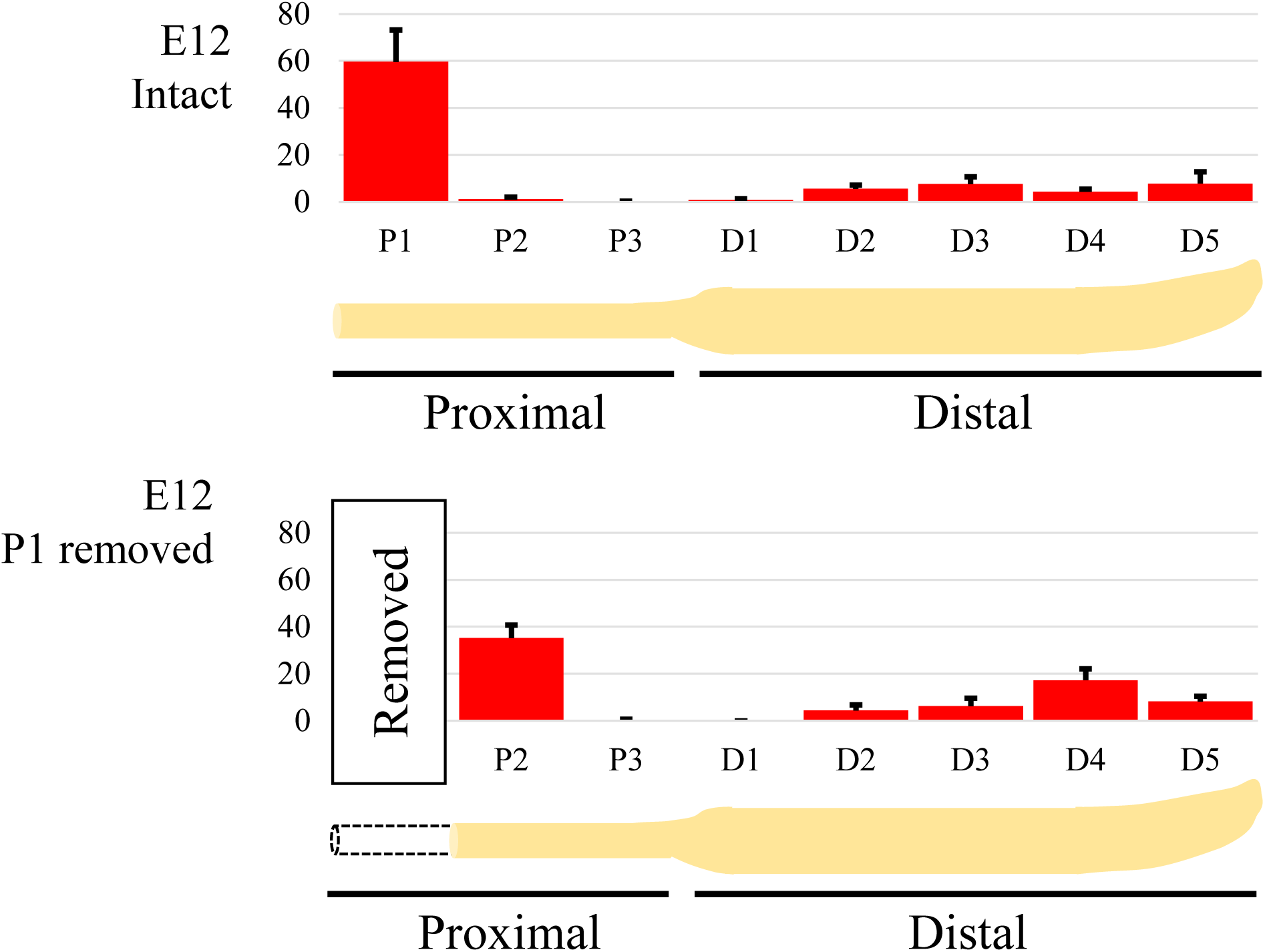
P1 removal increases OPW frequency in the adjacent P2 region. In the upper panel representing OPW distribution in intact E12 caecum, the P1 region of the E12 caecum exhibits a significantly higher frequency of OPW generation (the same graph as Fig. 1D, E12, n = 5), while the neighboring P2 region shows a significantly lower frequency. Removal of the P1 region results in an increased frequency of OPWs in the remaining P2 region (n = 5). Removal analysis was performed in the condition of short-term ex-vivo culture (Materials and methods).

**Figure S2.**
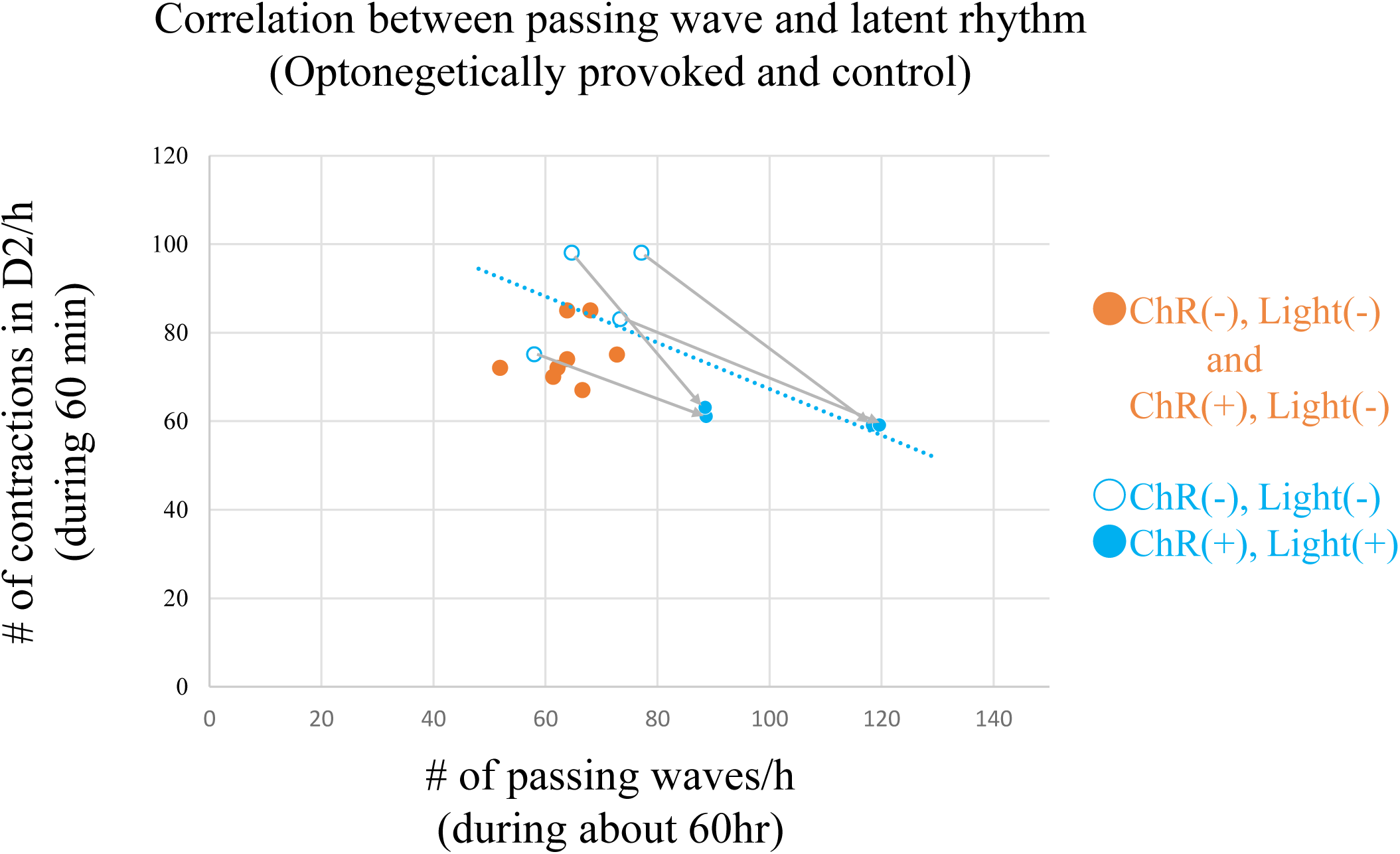
Optogenetically-increased frequency of passing waves during development reduces latent rhythms. Paired scatter plots showing relationship between the number of passing waves generated by optogenetic stimulations (Fig 4E, left panel) and the acquired number of contractions in the D2 region (Fig 4E, right panel) by blue dots. Blue open circles represent not-electroporated samples, while blue filled circles represent electroporated and irradiated samples, regression line is shown as blue dotted line. Values in no-blue light condition (Fig 4D) are plotted with orange dots.

## Supplementary movies

**Movie S1**

**Peristaltic movements in an intact E10 caecum related to Fig. 1C**

Time-lapse imaging of an intact E10 caecum under short-term ex vivo culture conditions (Materials and methods). Red circles indicate OPW generation sites. The dotted line separates the proximal and distal regions. Time is indicated in minutes:seconds.

**Movie S2**

**Peristaltic movements in an intact E12 caecum related to Fig. 1C**

Time-lapse imaging of an intact E12 caecum under short-term ex vivo culture conditions (Materials and methods). Red circles indicate OPW generation sites. The dotted line separates the proximal and distal regions. Time is indicated in minutes:seconds.

**Movie S3**

**Active contractions of caecal fragment related to Fig. 1D**

Time-lapse imaging of a fragmented caecum in a PDMS canal under short-term ex vivo culture conditions (Materials and methods). Time is indicated in minutes:seconds.

**Movie S4**

**Resetting of latent rhythm by mechanical stimulation related to Fig. 2.**

Time-lapse imaging of a P2 caecum fragment under short-term ex vivo culture conditions (Materials and methods). Upper panel: Spontaneous (spont.) and mechanically induced (induced) contractions are shown. Lower panel: A timeline indicating the timing of spontaneous and induced contractions. Time is indicated in minutes:seconds.

**Movie S5**

**Optogenetic inductions of peristaltic waves related to Fig. 4C.**

Time-lapse imaging of an E10 caecum expressing ChR2(D156C) under long-term ex vivo culture conditions (Materials and methods). Control caecum (ChR-) and optogenetically manipulated caecum (ChR+) were mounted on the PDMS canal. Left: proximal; Right: distal direction. Blue light pulses (30-second intervals) were applied to the ChR+ caecum to induce peristaltic waves, which propagated through the D2 region. Time is indicated in minutes:seconds.

## Notes

### Competing Interest Statement

The authors have declared no competing interest.

